# Accelerated dimensionality reduction of single-cell RNA sequencing data with fastglmpca

**DOI:** 10.1101/2024.03.23.586420

**Authors:** Eric Weine, Peter Carbonetto, Matthew Stephens

## Abstract

**Summary:** Motivated by theoretical and practical issues that arise when applying Principal Components Analysis (PCA) to count data, Townes et al introduced “Poisson GLM-PCA”, a variation of PCA adapted to count data, as a tool for dimensionality reduction of single-cell RNA sequencing (RNA-seq) data. However, fitting GLM-PCA is computationally challenging. Here we study this problem, and show that a simple algorithm, which we call “Alternating Poisson Regression” (APR), produces better quality fits, and in less time, than existing algorithms. APR is also memory-efficient, and lends itself to parallel implementation on multi-core processors, both of which are helpful for handling large single-cell RNA-seq data sets. We illustrate the benefits of this approach in two published single-cell RNA-seq data sets. The new algorithms are implemented in an R package, fastglmpca.

**Availability and implementation:** The fastglmpca R package is released on CRAN for Windows, macOS and Linux, and the source code is available at github.com/stephenslab/fastglmpca under the open source GPL-3 license. Scripts to reproduce the results in this paper are also available in the GitHub repository.

**Supplementary information:** Supplementary data are available on *BioRxiv* online.

---

Almost every analysis of single-cell RNA sequencing (scRNA-seq) data involves some kind of dimensionality reduction to help summarize and denoise the data (Amezquita et al., 2020; Linderman, 2021; Stuart et al., 2019; Sun et al., 2019; Tsuyuzaki et al., 2020). Principal Components Analysis (PCA) is a widely used dimensionality reduction technique, but it has been criticized as being poorly suited to the sparse count nature of scRNA-seq data. Motivated by this, Townes et al. (2019) suggested instead using a version of PCA, called “GLM-PCA”, that is specifically tailored to count data. However, GLM-PCA is computationally challenging to fit. In this paper we provide faster algorithms, implemented in the software `fastglmpca`, to fit this model.

The GLM-PCA model combines PCA with ideas from generalized linear models (McCullagh, 1989) and dates back at least to Choulakian (1996); see also Collins et al. (2001) and Chen et al. (2013). We consider here the Poisson version of this model which was the primary focus in Townes et al. (2019). The Poisson variant models the *n × m* data matrix **Y** as

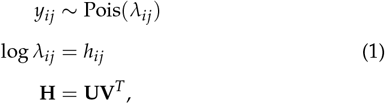

where Pois(*λ*) denotes the Poisson distribution with mean *λ*; *y*_*ij*_ and *h*_*ij*_ denote entries of the matrices **Y** and **H**, respectively; **U** ∈ **R**^*n×K*^ and **V** *∈* **R**^*m×K*^ are the matrices of unknowns to be estimated from the data; and *K >* 0 is an integer specifying the dimension of the reduced representation, typically a number much smaller than *n* or *m*. In this form, the model is symmetric in the rows and columns of **Y**, but by convention we assume that rows *i* are genes and columns *j* are cells (e.g., Nicol and Miller, 2023; Stuart et al., 2019; Townes et al., 2019). See the Supplementary Text for elaborations of this model with options to specify row (gene) and column (cell) covariates.

In (1), we do not impose the typical PCA constraints enforcing orthogonality on **U** and **V** because such constraints can be easily applied after fitting; that is, once **U, V** have been estimated, a “PCA-like” decomposition for **H** can be obtained from a singular value decomposition of the estimated **UV**^*T*^. See the Supplementary Text for details.

Whereas standard PCA involves straightforward application of a (truncated) SVD algorithm, fitting the GLM-PCA model is much less straightforward; computing a maximum-likelihood estimate (MLE) of **U, V** in (1) is a high-dimensional, nonconvex optimization problem. The `glmpca` R package (Townes et al., 2019) uses *stochastic gradients*, progressively improving the parameter estimates in the direction of a noisy estimate of the gradient computed with a random subset (a “mini-batch”) of the cells. (The R package `NewWave` afits a related model via stochastic gradients; see Agostinis et al. 2022.) Since the performance of the stochastic gradients method can depend strongly on the choice of learning rate, they used the adaptive AvaGrad method (Savarese et al., 2021). However, even with AvaGrad, the method can be very sensitive to the choice of learning rate, and may be unstable if the learning rate is too large. (`glmpca` implements other approaches, but we and others Nicol and Miller 2023 have found that the AvaGrad approach generally performed best.)

## Algorithm 1 Sketch of Alternating Poisson Regression for GLM-PCA. Row *i* and column *j* of Y are denoted, respectively, by *y*_:,*i*_ and *y*_:,*j*_.

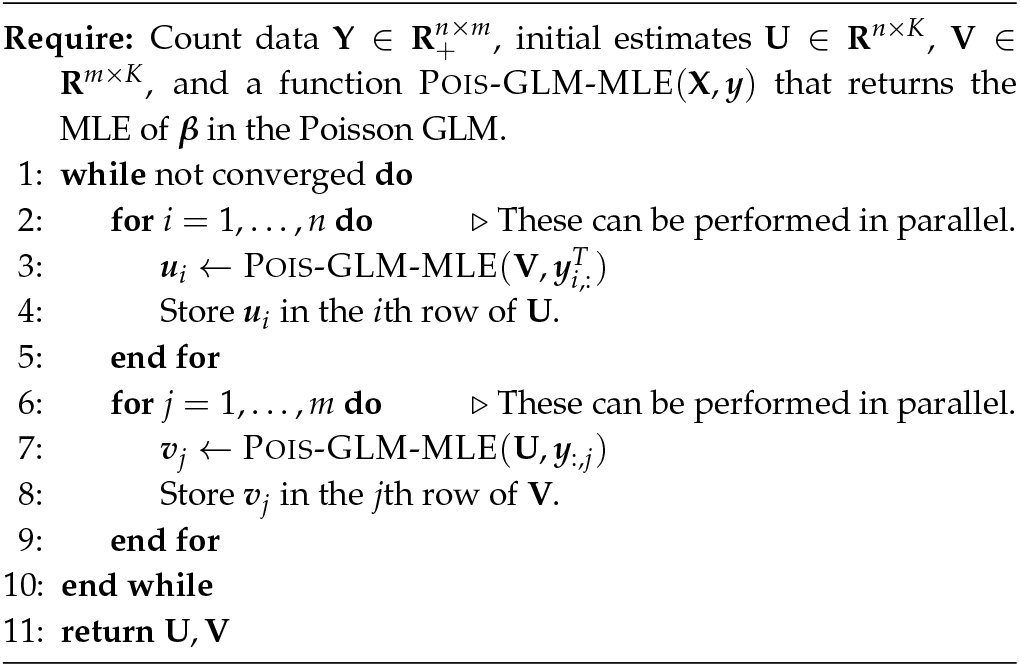

The `scGBM` R package (Nicol and Miller, 2023) takes a different approach, iteratively solving an approximation to the log-likelihood that has the form of a more tractable “weighted SVD” problem. This approach, called IRSVD (“iteratively reweighted SVD”), can be very memory-intensive—for example, it involves forming a matrix of the same size as **Y** that is not sparse—which limits its application to larger scRNA-seq data sets (see also Lee and Han 2022 for related discussion of these issues).

Here we describe another approach to fitting GLM-PCA models that is based on a simple observation: when **V** is fixed, computing the MLE for **U** reduces to the much simpler and very well studied problem of independently fitting *n* generalized linear models (GLMs) with a Poisson error distribution and log-link function (McCullagh, 1989). Similarly, when **U** is fixed, computing the MLE of **V** reduces to independently fitting *m* Poisson GLMs. This suggests a *block-coordinate optimization approach* (Wright, 2015) that alternates between optimizing **U** with fixed **V**, and optimizing **V** with fixed **U** (Algorithm 1). This approach is analogous to the “alternating least squares” algorithm for truncated SVD (Hastie et al., 2015), and a similar alternating approach has proven very effective for non-negative matrix factorization. We call our approach “Alternating Poisson Regression” (APR) to draw attention to its two key aspects:

(i) the alternating optimization of **U** and **V**, and (ii) the reduction to simpler Poisson GLM optimization problems. We have implemented the APR algorithm in the R package `fastglmpca`.

The APR approach has several benefits. First, it has strong convergence guarantees; the block-coordinatewise updates monotonically improve the log-likelihood, and under mild conditions converge to a (local) maximum of the likelihood (Wright, 2015). In addition, by splitting the large optimization problem into smaller pieces (the Poisson GLMs), the computations are memory-efficient and are trivially parallelized to take advantage of multi-core processors.

Since APR reduces the problem of fitting a Poisson GLM-PCA model to the problem of fitting many (much smaller) Poisson GLMs, the speed of the APR algorithm depends critically on how efficiently one can fit the individual Poisson GLMs. The “classic” algorithm for GLMs is iteratively reweighted least squares (IRLS) (McCullagh, 1989). However, the complexity of IRLS grows very quickly with *K*— we would prefer an approach that is still fast for large *K*. Therefore, we instead take a cyclic coordinate descent (CCD) approach to fitting each Poisson GLM, which involves very simple (and therefore very fast) 1-d Newton updates of the GLM parameters. Although the convergence behaviour of CCD is theoretically much worse than IRLS, the CCD updates can converge quickly in practice, especially when we orthogonalize **U** and **V** at each iteration. These and other details of the implementation resulting in improved speed and reliability are discussed in the Supplementary Text.

To illustrate the benefits of the APR approach for GLM-PCA, we analyzed three scRNA-seq data sets: 7,193 cells from the tracheal epithelium in wild-type mice (Montoro et al., 2018); and two peripheral blood mononuclear cell (PBMC) data sets with 68,579 cells (“68k PBMC”) and 94,655 cells (“95k PBMC”) (Zheng et al., 2017). We compared APR, implemented in the R package `fastglmpca`, and two existing software implementations (also in R): the Iterative Reweighted SVD (IRSVD) algorithm implemented in the R package `scGBM` (Miller and Carter, 2020; Nicol and Miller, 2023); and the adaptive stochastic gradient algorithm (“AvaGrad”) implemented in the R package `glmpca` (Savarese et al., 2021; Townes et al., 2019).

Since all the methods optimize the same objective function (the log-likelihood), we used this objective function to compare the quality of the fits. The quality of the fit and the running time depends very strongly on the criterion used to stop the model fitting. Since this criterion is somewhat arbitrary, we performed these comparisons by visualizing the evolution of the log-likelihood against running time. The results for different settings of *K*, ranging from 2 to 25, are summarized in Supplementary Figures 1 and 2, and for *K* = 10 in Fig. 1A. (Full details of these comparisons are given in the Supplementary Text.) The typical result was that while all the algorithms continued to (slowly) improve the fits even after running for many hours, the APR algorithm improved the fit at a much greater rate than the other approaches. (In two exceptions to this, AvaGrad seemed to have settled into better local solutions than APR; Supplementary Fig. 1.) Examining the log-likelihood for each cell reveals that `fastglmpca` consistently improved the fit across almost all cells, rather than just a small subset of cells (Supplementary Figures 3, 4). When multi-processor computing resources were available, APR leveraged these resources to dramatically speed up model fitting. For example, to achieve the same log-likelihood as running AvaGrad for 10 hours, the parallel APR updates running on 28-core processor needed to run only 10 minutes on the 68k PBMC data set and only 1 minute on the epithelial airway data set (Fig. 1A).

**Fig. 1:**
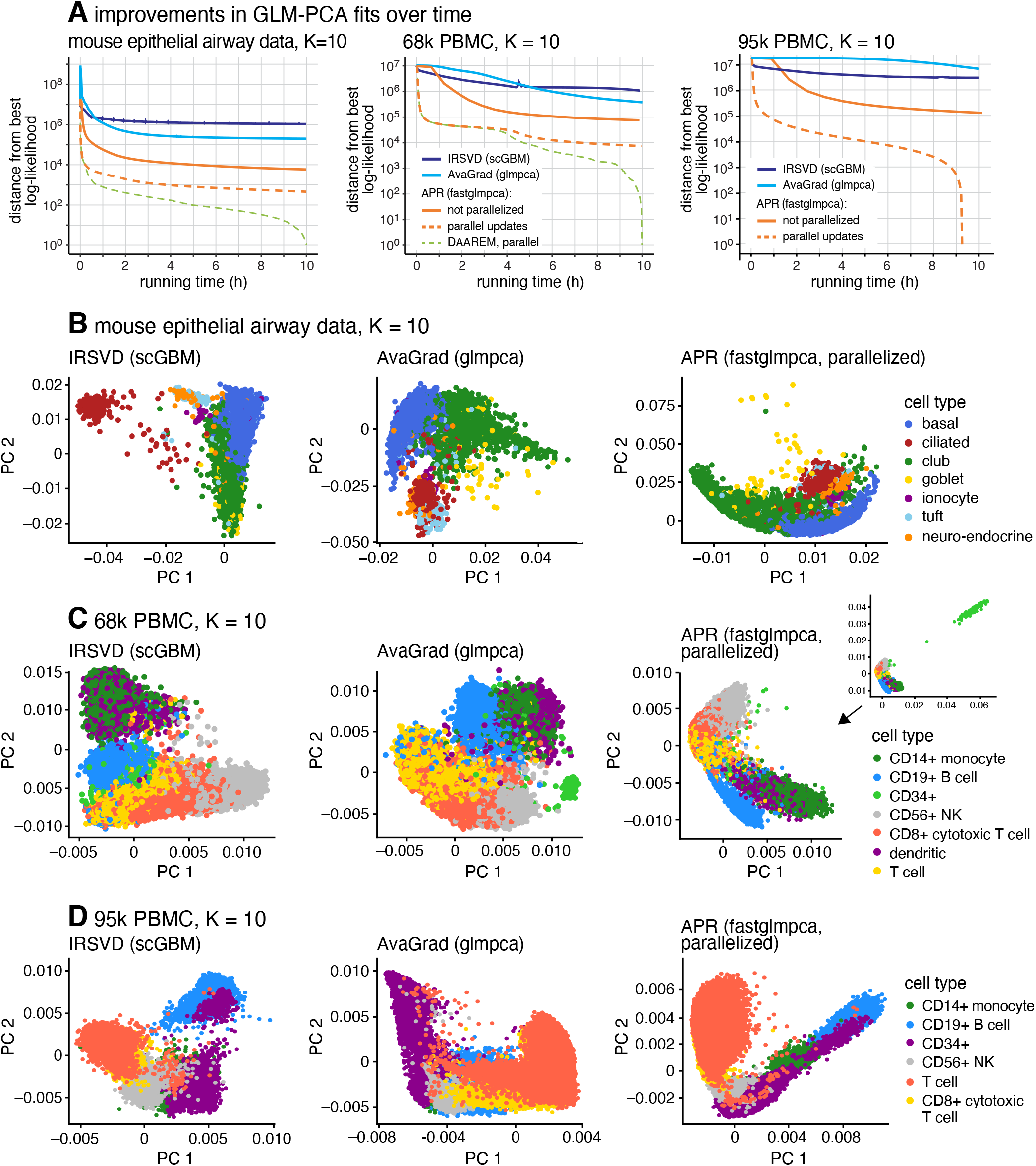
Comparison of GLM-PCA model fitting algorithms on three scRNA-seq data sets: mouse epithelial airway data (7,193 cells), 68k PBMC data (68,579 cells) and 95k PBMC data (94,655 cells). **(A)** Improvement in *K* = 10 GLM-PCA fits over time. Log-likelihoods are shown relative to the best log-likelihood recovered among methods compared. The Y axis has a log scale, and log-likelihood differences less than 1 are shown as 1. **(B–D)** The first 2 rows of **V** after fitting the model for about 10 hours. The cell types in B and C are estimates from scRNA-seq data (Montoro et al., 2018; Zheng et al., 2017); the cell types in D are the sorted cell populations (Zheng et al., 2017).

To assess whether the different log-likelihoods achieved by different methods corresponded to qualitatively different representations of the data, we examined the estimated latent factors (the “PCs”) returned by each method. We found that the different methods sometimes generated quite different representations; consider for example the results with *K* = 10 shown in Fig. 1B–D and in Supplementary Figures 5–7). Since there is no ground truth, we cannot definitively say whether one representation is better than the other. (Overall, the APR estimates seemed to produce more striking visual representations of the key cell population substructures, but this is subjective.) The 95k PBMC data set does contain an independent ground truth of sorts since it is made up of sorted cell populations (Fig. 1D; Supplementary Text). Therefore, we performed a downstream clustering analysis on each estimate of **V**, and then, following Sun et al. (2019), used two measures – normalized mutual information (NMI) and adjusted rand index (ARI) – to assess how well the clusters recovered the sorted cell types (Supplementary Fig. 8). The (parallelized) APR updates very quickly produced clusters that closely recovered the sorted cell populations, whereas AvaGrad took much longer to obtain clusters of similar accuracy.

IRSVD also produced reasonably good clusters quickly, but then seemed to get “stuck” and had trouble making further improvements.

All algorithms have a computational expense that grows at most linearly in *n, m* and *K* (Supplementary Fig. 9). However, `fastglmpca` can cope with much larger scRNA-seq data sets because of the care taken to avoid computations that “fill in” the sparse data matrix **Y**. In summary, the key differentiating factors are (1) the speed at which the `fastglmpca` updates find good GLM-PCA fits, particularly when the updates can be run in parallel, and (2) numerical computations that greatly reduce memory usage when the data matrix **Y** is large and sparse. We also observed, anecdotally, the potential to additionally accelerate model fits using DAAREM (Henderson and Varadhan, 2019) (Fig. 1A).

The Poisson GLM-PCA model (1) can be seen as combining a Poisson measurement model with a low-rank (log) expression model (in the terminology of Sarkar and Stephens 2021). There is good theoretical and empirical support for the Poisson measurement model, but the expression model would likely be improved by allowing for deviation from an exact low-rank structure (Sarkar and Stephens, 2021). This idea can motivate alternative models, such as the negative binomial variation of GLM-PCA that is implemented in `glmpca` (see also the `NewWave` package; Agostinis et al. 2022.) Future work could consider extending the algorithms introduced here to the negative binomial case.

In summary, we contribute a new R package, `fastglmpca`, which implements fast algorithms for dimensionality reduction of count data based on the Poisson GLM-PCA model. The package is available on CRAN for all major computing platforms. It features a well-documented, user-friendly model fitting interface that aligns closely with `glmpca` and `scGBM`, and a vignette giving an example analysis of scRNA-seq data. The interface splits the GLM-PCA analysis into two phases: an initialization phase, where modeling choices are made, including the rank, *K*, and row- and column-covariates (function “`init glmpca pois`”); and a model fitting phase, where the optimization may be monitored and fine-tuned (function “`fit glmpca pois`”). The core model fitting routines were implemented efficiently in C++ using the Armadillo linear algebra library and Intel Threading Building Blocks (TBB).

## Acknowledgments

We thank the Research Computing Center for providing high-performance computing resources used in this research. We thank Abhishek Sarkar, Jason Willwerscheid, Dongyue Xie and other members of the Stephens lab for helpful suggestions and feedback.

