## Supplementary information for "Accelerated dimensionality reduction of single-cell RNA sequencing data with fastglmpca"

### SUPPLEMENTARY MATERIALS: RAPID DIMENSIONALITY REDUCTION OF SINGLE-CELL RNA SEQUENCING DATA WITH FASTGLMPCA

**1. General problem formulation.** Let $\mathbf{Y} \in \mathbf{R}_+^{n \times m}$ be an $n \times m$ non-negative matrix storing observed counts $y_{ij}$. Similar to [9, 20, 27, 28], we consider the following model of the count data:

$$
\begin{aligned}
y_{ij} &\sim \text{Pois}(\lambda_{ij}) \\
\log \lambda_{ij} &= h_{ij} \\
\mathbf{H} &= \mathbf{U\Sigma V}^T + \mathbf{XB}^T + \mathbf{WZ}^T,
\end{aligned}
\tag{1}
$$

in which Pois($\lambda$) denotes the Poisson probability distribution with mean $\lambda$, $\mathbf{X} \in \mathbf{R}^{n \times n_x}, \mathbf{Z} \in \mathbf{R}^{m \times n_z}$ are additional observed covariates for, respectively, the rows and columns of $\mathbf{Y}$ ($n_x$ or $n_z$, or both, may be zero), $\mathbf{U} \in \mathbf{R}^{n \times K}$, $\mathbf{\Sigma} \in \mathbf{R}^{K \times K}$, $\mathbf{V} \in \mathbf{R}^{m \times K}$, $\mathbf{B} \in \mathbf{R}^{m \times n_x}$, $\mathbf{W} \in \mathbf{R}^{n \times n_z}$ are matrices of unknowns to be estimated from the data, and $K > 0$ is an integer specifying the rank of the matrix factorization $\mathbf{U\Sigma V}^T$, which is typically much smaller than $n$ or $m$. (Our software implementation also allows for fixing one or more columns of $\mathbf{B}$ and/or one or more columns of $\mathbf{W}$ to chosen values.)

In the simplest case when there are no additional covariates, $n_x = n_z = 0$, the model is

$$
\begin{aligned}
y_{ij} &\sim \text{Pois}(\lambda_{ij}) \\
\log \lambda_{ij} &= h_{ij} \\
\mathbf{H} &= \mathbf{U\Sigma V}^T,
\end{aligned}
\tag{2}
$$

To make the unknowns identifiable (up to sign) under general conditions [20], we further impose the following "PCA-like" constraints:

$$
\begin{aligned}
\mathbf{U}^T\mathbf{U} &= \mathbf{I}_K \\
\mathbf{V}^T\mathbf{V} &= \mathbf{I}_K \\
\mathbf{\Sigma} &= \text{diag}(\sigma_1, \ldots, \sigma_K), \quad \infty > \sigma_1 > \cdots > \sigma_K > 0,
\end{aligned}
\tag{3}
$$

in which $\mathbf{I}_n$ denotes the $n \times n$ identity matrix. We call these the "identifiability constraints."

Our aim is to compute maximum-likelihood estimates of the unknowns $\mathbf{U}$, $\mathbf{\Sigma}$, $\mathbf{V}$, $\mathbf{B}$, and $\mathbf{W}$ in (1) subject to the identifiability constraints (3). As we will show, computing the constrained maximum-likelihood solution for (2) is easily extended to the model with additional covariates (1), so we focus initially on fitting (2), then we explain the extension to (1).

*Laboratory for Information and Decision Systems, Massachusetts Institute of Technology, Cambridge, MA, USA.
†Department of Human Genetics, University of Chicago, Chicago, IL, USA.
‡Departments of Statistics and Human Genetics, University of Chicago, Chicago, IL, USA

**1.1. Dealing with the identifiability constraints.** To deal with the identifiability constraints (3), we first compute maximum-likelihood estimates of $\mathbf{U}$ and $\mathbf{V}$ without these constraints, then we transform the solution *post hoc* to find equivalent estimates of $\mathbf{U}$, $\mathbf{V}$, and $\mathbf{\Sigma}$ satisfying the identifiability constraints.

Denote the unconstrained estimates by $\hat{\mathbf{U}}$ and $\hat{\mathbf{V}}$. Then a solution $\mathbf{U}, \mathbf{\Sigma}, \mathbf{V}$ satisfying the constraints (3) is easily recovered by forming $\hat{\mathbf{H}} = \hat{\mathbf{U}}\hat{\mathbf{V}}^T$, then obtaining the truncated rank-$K$ singular value decomposition (SVD),

$$\mathbf{U}, \mathbf{\Sigma}, \mathbf{V} \leftarrow \mathrm{SVD}(\hat{\mathbf{H}}). \tag{4}$$

The solutions $\hat{\mathbf{U}}, \hat{\mathbf{V}}$ and $\mathbf{U}, \mathbf{\Sigma}, \mathbf{V}$ have the same likelihood because $\hat{\mathbf{H}} = \hat{\mathbf{U}}\hat{\mathbf{V}}^T = \mathbf{U}\mathbf{\Sigma}\mathbf{V}^T$, and $\mathbf{U}, \mathbf{\Sigma}, \mathbf{V}$ satisfy the identifiability constraints (3) by construction. However, we would like to avoid computing the SVD of an $n \times m$ matrix, which can be computationally expensive and memory-intensive. The following procedure generates the same result as (4), but avoids the costly SVD:

$$\begin{aligned}
\mathbf{Q}_U, \mathbf{R}_U &\leftarrow \mathrm{QR}(\hat{\mathbf{U}}) \\
\mathbf{Q}_V, \mathbf{R}_V &\leftarrow \mathrm{QR}(\hat{\mathbf{V}}) \\
\tilde{\mathbf{U}}, \mathbf{\Sigma}, \tilde{\mathbf{V}} &\leftarrow \mathrm{SVD}(\mathbf{R}_U \mathbf{R}_V^T) \\
\mathbf{U} &\leftarrow \mathbf{Q}_U \tilde{\mathbf{U}} \\
\mathbf{V} &\leftarrow \mathbf{Q}_V \tilde{\mathbf{V}}
\end{aligned} \tag{5}$$

The first two steps compute QR decompositions of $\hat{\mathbf{U}}$ and $\hat{\mathbf{V}}$. Since $\mathbf{R}_U$ and $\mathbf{R}_V$ are $K \times K$ matrices, the product of these two matrices is very quickly computed. Therefore, the predominant computation in this procedure are the QR decompositions, each of which have a much more favorable time complexity than the truncated SVD of $\hat{\mathbf{H}}$.

**1.2. Unconstrained optimization problem.** Since the identifiability constraints are easily satisfied *post hoc*, for the remainder we focus on developing algorithms for the unconstrained optimization of $\mathbf{U}$ and $\mathbf{V}$. Arbitrarily setting $\mathbf{\Sigma} = \mathbf{I}_K$, we seek to maximize the following objective, which is the log-likelihood derived from model (2) after removing terms that do not involve $\mathbf{U}$ or $\mathbf{V}$:

$$\begin{aligned}
\ell_{\text{GLM-PCA}}(\mathbf{Y}; \mathbf{U}, \mathbf{V}) &= \sum_{i=1}^{n}\sum_{j=1}^{m} y_{ij} \log \lambda_{ij} - \sum_{i=1}^{n}\sum_{j=1}^{m} \lambda_{ij} \\
&= \sum_{i=1}^{n}\sum_{j=1}^{m}\sum_{k=1}^{K} y_{ij} u_{ik} v_{jk} - \sum_{i=1}^{n}\sum_{j=1}^{m} \exp\left(\sum_{k=1}^{K} u_{ik} v_{jk}\right).
\end{aligned} \tag{6}$$

In the next section, we develop a fast algorithm for finding an unconstrained maximizer of (6).

**2. Algorithm for maximizing the GLM-PCA log-likelihood.** To approach the problem of finding an unconstrained maximizer of the GLM-PCA log-likelihood (6), we first consider the much simpler problem of fitting the following model to data $\mathbf{X} \in \mathbf{R}^{N \times K}$, $\boldsymbol{y} \in \mathbf{R}^N$:

$$\begin{aligned}
y_i &\sim \mathrm{Pois}(\lambda_i) \\
\log \lambda_i &= h_i \\
\boldsymbol{h} &= \mathbf{X}\boldsymbol{\beta},
\end{aligned} \tag{7}$$

in which $\boldsymbol{\beta} = (\beta_1, \ldots, \beta_K) \in \mathbf{R}^K$ is the vector of $K$ unknowns to be estimated from the data $\mathbf{X}, \boldsymbol{y}$. This is the standard generalized linear model (GLM) with a Poisson error distribution and log-link function [16]. Therefore, we refer to (7) as the "Poisson GLM." For later convenience, we define a function that returns the maximum-likelihood estimate (MLE) of the coefficients $\boldsymbol{\beta}$ in the Poisson GLM:

$$\text{POIS-GLM-MLE}(\mathbf{X}, \boldsymbol{y}) := \underset{\boldsymbol{\beta} \in \mathbf{R}^K}{\operatorname{argmax}} \, \ell_{\text{POIS-GLM}}(\mathbf{X}, \boldsymbol{y}; \boldsymbol{\beta}), \tag{8}$$

where $\ell_{\text{POIS-GLM}}(\mathbf{X}, \boldsymbol{y}; \boldsymbol{\beta})$ denotes the likelihood of the Poisson GLM (7).

We now relate the problem of computing the MLE for the Poisson GLM to our original problem. The likelihood for the Poisson GLM (7), excluding terms that do not involve $\boldsymbol{\beta}$, is

$$\begin{aligned} \ell_{\text{POIS-GLM}}(\mathbf{X}, \boldsymbol{y}; \boldsymbol{\beta}) &= \sum_{i=1}^{N} y_i \log \lambda_i - \sum_{i=1}^{N} \lambda_i \\ &= \sum_{i=1}^{N} \sum_{k=1}^{K} x_{ik} y_i \beta_k - \sum_{i=1}^{N} \exp\left( \sum_{k=1}^{K} x_{ik} \beta_k \right). \end{aligned} \tag{9}$$

By rearranging (6), the GLM-PCA log-likelihood can be rewritten as a sum of Poisson GLM log-likelihoods in two different ways:

$$\ell_{\text{GLM-PCA}}(\mathbf{Y}; \mathbf{U}, \mathbf{V}) = \sum_{i=1}^{n} \ell_{\text{POIS-GLM}}(\boldsymbol{y}_{i,:}^T, \mathbf{V}; \boldsymbol{u}_{i,:}^T) \tag{10}$$

$$= \sum_{j=1}^{m} \ell_{\text{POIS-GLM}}(\boldsymbol{y}_{:,j}, \mathbf{U}; \boldsymbol{v}_{j,:}^T), \tag{11}$$

in which $\boldsymbol{y}_{i,:} \in \mathbf{R}_+^m$, $\boldsymbol{y}_{:,j} \in \mathbf{R}_+^n$ denote vectors storing, respectively, the $i$th row and $j$ column of $\mathbf{Y}$, and $\boldsymbol{u}_{i,:} \in \mathbf{R}^K, \boldsymbol{v}_{j,:} \in \mathbf{R}^K$ denote vectors storing, respectively, the $i$th row of $\mathbf{U}$ and the $j$th row of $\mathbf{V}$. This result, while straightforward algebraically, has an important implication: if we fix $\mathbf{V}$, the GLM-PCA optimization problem reduces to a collection of $n$ Poisson GLM optimization problems (eq. 10) and, similarly, if we fix $\mathbf{U}$, the GLM-PCA optimization problem reduces to a collection of $m$ Poisson GLM optimization problems (eq. 11). This suggests an *alternating optimization* approach in which we iterate between optimizing $\mathbf{U}$ with $\mathbf{V}$ fixed, and optimizing $\mathbf{V}$ with $\mathbf{U}$ fixed. This is an example of a block-coordinate optimization algorithm [33] (also "nonlinear Gauss-Seidel" [5]), where the "blocks" are $\mathbf{U}$ and $\mathbf{V}$. It is analogous to "alternating least squares" for matrix factorization with Gaussian errors, and therefore we call this algorithm "Alternating Poisson Regression" (APR). (Similarly, in non-negative matrix factorization some algorithms reduce to solving a series of non-negative least squares problems [7, 12, 14, 15].) This idea is summarized by Algorithm 1 in the main text.

The core computation of the APR algorithm is the MLE of the Poisson GLM. Therefore, in the next section we focus on the development of a fast solution to this core computation.

**2.1. Fitting the Poisson GLM.** We compute the MLE for the Poisson GLM (7) using a *cyclic coordinate descent* (CCD) algorithm [5, 33], in which each coordinate $\beta_k$ is optimized one at a time while the others are fixed. While the iteratively reweighted least squares (IRLS) algorithm is the standard algorithm for fitting GLMs, including the Poisson GLM, the complexity of IRLS is $O(K^3)$ due to the inversion or factorization of a $K \times K$ matrix. To handle large data sets, we would prefer a more efficient alternative that scales better to larger values of $K$. For this reason, we use the CCD approach since its computational complexity grows only linearly with $K$.

The CCD algorithm performs the following 1-d optimization for each $k = 1, \ldots, K$,

$$\beta_k^{\text{new}} \leftarrow \text{argmax}_{\beta_k \in \mathbf{R}} \, \ell_{\text{POIS-GLM}}(\mathbf{X}, \boldsymbol{y}; \boldsymbol{\beta}), \tag{12}$$

and repeats these 1-d optimizations until the iterates $\boldsymbol{\beta}^{(0)}, \boldsymbol{\beta}^{(1)}, \ldots$ reach a stationary point. (Although we have framed this as a maximum-likelihood estimation problem, the convention in most numerical optimization papers is to frame the optimization as as a minimization problem, so we view our algorithms as minimizing the *negative log-likelihood*, hence "descent.")

Note that this coordinate descent algorithm is different from the coordinate descent approach for GLMs in `glmnet` [11]; `glmnet` essentially implements IRLS, but performs coordinatewise updates to solve each weighted least-squares problem, whereas we perform the coordinatewise updates (12) on the original log-likelihood. For example, our approach is similar to the cyclic coordinate ascent algorithm for penalized logistic regression described in [34].

To further motivate the CCD algorithm, consider the first and second-order derivatives of Poisson GLM log-likelihood (9):

$$g_k := \frac{\partial \ell_{\text{POIS-GLM}}}{\partial \beta_k} = \sum_{i=1}^{N} x_{ik}(y_i - \lambda_i) \tag{13}$$

$$h_{kk'} := \frac{\partial^2 \ell_{\text{POIS-GLM}}}{\partial \beta_k \partial \beta_{k'}} = -\sum_{i=1}^{N} x_{ik} x_{ik'} \lambda_i. \tag{14}$$

We showed previously that it is possible to easily orthogonalize $\mathbf{U}$ and $\mathbf{V}$. When solving Pois-GLM-MLE$(\mathbf{X}, \boldsymbol{y})$, $\mathbf{X}$ will either be $\mathbf{U}$ or $\mathbf{V}$, so we can assume that $\mathbf{X}$ is orthogonal simply by making it so. Although the $K \times K$ matrix of second-order partial derivatives (the Hessian) is not guaranteed to be diagonal even when $\mathbf{X}$ is orthogonal, it might be close to diagonal; that is, the diagonal elements may be much larger than the non-diagonal elements. In the special case in which all the $\lambda_i$'s are the same, the off-diagonal entries of the Hessian (eq. 14 with $k \neq k'$) will be zero. Therefore, in situations where the $\lambda_i$'s are similar, the Hessian should be close to diagonal. Coordinate descent algorithms have good rates of convergence when the Hessian is diagonal or close to diagonal. Even when the Hessian is not close to diagonal, in practice coordinatewise algorithms often converge very rapidly. In summary, although the worst-case convergence behaviour of (cyclic) coordinate descent is poor, we view CCD as a practical approach that it very scalable and should have good convergence behaviour in most situations.

**2.2. Other implementation details.** Here we describe other important aspects of the computer implementation.

**2.2.1. Fitting the Poisson GLMs in parallel.** The $n$ Poisson GLM optimizations in (10) and the $m$ Poisson GLM optimizations in (11) are trivially parallelizable, so we can take advantage of multicore processors, for example, to greatly speed up the optimization. The core model fitting routines were implemented efficiently in C++ using the Armadillo linear algebra library. And we used Intel Threading Building Blocks (TBB) multithreading [10], interfaced via the RcppParallel R package [2], to solve the Poisson GLM subproblems in parallel.

**2.2.2. Initialization.** To initialize the model fitting, by default we set the entries of $\mathbf{U}$ and $\mathbf{V}$ to random values (drawn from a normal distribution). While other approaches have been suggested (e.g., [20]), we find that these initializations did not reliably lead to better solutions nor faster convergence.

**2.2.3. Stopping criteria.** We stop the model fitting when a maximum number of outer-loop iterations in Algorithm 1 is reached (chosen by the user), or when two successive outer-loop iterates improve the log-likelihood (6) by less than some tolerance, $\varepsilon > 0$. This tolerance value is also set by the user. By default, $\varepsilon = 10^{-4}$.

**2.2.4. CCD algorithm.** The CCD updates (12) were implemented using backtracking line search [21, Algorithm 3.1], with Newton search direction $p_k = g_k/h_{kk}$. While there is some debate as to whether line search is worth the added expense (e.g., [11]), in this setting we have found it to be important for avoiding diverging updates.

Fully optimizing $\mathbf{U}$ with fixed $\mathbf{V}$ (and fully optimizing $\mathbf{V}$ with fixed $\mathbf{U}$) is possible (up to some numerical error), but in practice it is not necessary; because of the alternating nature of the algorithm, incompletely optimizing $\mathbf{U}$ and $\mathbf{V}$ provides considerable reductions in computation with little impact on convergence. Specifically, we approximated (12) with the solution obtained by cycling through the CCD updates a fixed number of times, in which $\boldsymbol{\beta}$ was initialized to row $i$ of $\mathbf{U}$ or row $j$ of $\mathbf{V}$. The number of cycles to perform in CCD is a tuning parameter in the software. We found that cycling through coordinates 1 through $K$ three times worked well in practice, and so we took this to be the software default. Although the theory on coordinate descent does not guarantee convergence unless we compute exact solutions, we have not encountered situations in which inexact solutions caused the iterates to diverge.

**2.2.5. Acceleration of the outer-loop updates.** Since APR is a coordinatewise optimization method that alternates between optimizing $\mathbf{U}$ and optimizing $\mathbf{V}$, one possible reason for slow convergence is strong interdependence between $\mathbf{U}$ and $\mathbf{V}$ [4, 33]. To remedy this issue, we tried DAAREM, an off-the-shelf method for accelerating fixed-point iterations [13, 26]. The DAAREM-accelerated variant of APR is the same as the original algorithm, except for an additional step at the end of each iteration of the outer-loop that computes a modified update. The additional computation involved in this additional step is negligible compared to the other computations.

We made two minor changes to the optimization which greatly improved the effectiveness of DAAREM for fitting GLM-PCA models. First, we found that DAAREM did not work well when $\mathbf{U}$ was made orthogonal before updating $\mathbf{V}$, and vice versa, so we skipped this orthogonalization step when using DAAREM.

Second, we found that the DAAREM-modified updates behaved poorly when the iterates were changing rapidly and were far away from the solution. Therefore, in our numerical experiments we first performed 100 unaccelerated APR updates to obtain a good initial estimate, then we performed a second round of APR updates with DAAREM.

**3. Comparisons in simulated data sets and scRNA-seq data sets.** The source code implementing these experiments is provided in the "inst" directory within the GitHub repository, `https://github.com/stephenslab/fastglmpca`. The Zenodo repository [32] contains a snapshot of this source code near the time of publication.

**3.1. Simulated data sets.** We simulated data matrices $\mathbf{Y}$ of different sizes to assess scalability of the model fitting algorithms. Specifically, for each trial, we simulated $\mathbf{Y}$ using the `generate_glmpca_data_pois()` function from the `fastglmpca` package. This function randomly generates a matrix $\mathbf{Y}$ from the GLM-PCA model (1) in which the parameters of the GLM-PCA model have been chosen to roughly mimic scRNA-seq data. The value of $K$ in the GLM-PCA model used to simulate the data was set to 5 or 10.

**3.2. Single-cell RNA-seq data sets.** All scRNA-seq data sets were stored as sparse $n \times m$ matrices $\mathbf{Y}$, where $n$ was the number of genes and $m$ was the number of cells.

Note that no additional steps were taken to preprocess the scRNA-seq UMI counts other than removing genes with no variation in the data. Since the GLM-PCA model is a Poisson model of UMI counts, no transformation or normalization of the counts is needed.

**3.2.1. Mouse epithelial airway data set.** The mouse epithelial airway data were gene expression profiles of trachea epithelial cells in C57BL/6 mice obtained using droplet-based 3' scRNA-seq, processed using the GemCode Single Cell Platform [17]. The data were downloaded from the Gene Expression Omnibus (GEO) website, accession GSE103354. Specifically, we downloaded file `GSE103354_Trachea_droplet_UMIcounts.txt.gz`. This file also contained predicted cell-type assignments from the community detection analysis of [17]. After removing genes that were not expressed in any of the cells, $\mathbf{Y}$ was a 18,388 $\times$ 7,193 matrix of UMI counts, and 90.7% of the counts were zero.

**3.2.2. 95k PBMC data set.** This data set combines reference transcriptome profiles generated from 10 bead-enriched subpopulations of PBMCs (Donor A) and processed using Cell Ranger 1.1.0 [35, 36]. To compile this data set, we downloaded the "Gene/cell matrix (filtered)" `tar.gz` file from the 10x Genomics website for each of the following 10 FACS-purified data sets: CD14+ monocytes, CD19+ B cells, CD34+ cells, CD4+ helper T Cells, CD4+/CD25+ regulatory T Cells, CD4+/CD45RA+/CD25- naive T cells, CD4+/CD45RO+ memory T Cells, CD56+ natural killer cells, CD8+ cytotoxic T cells and CD8+/CD45RA+ naive cytotoxic T cells. After combining these 10 data sets, then filtering out unexpressed genes, $\mathbf{Y}$ was a 21,952 $\times$ 94,655 matrix of UMI counts in which 97.1% of the counts were zero.

In Fig. 1 and Supplementary Fig. 7, the following cells were labeled as "T cells": CD4+ helper T Cells, CD4+/CD25+ regulatory T Cells, CD4+/CD45RA+/CD25- naive T cells, CD4+/CD45RO+ memory T Cells and CD8+/CD45RA+ naive cytotoxic T cells.

**3.2.3. 68k PBMC data set.** The 68k PBMC data were transcriptome profiles from peripheral blood mononuclear cells (PBMCs) processed using Cell Ranger 1.1.0 [35]. We downloaded the "Gene/cell matrix (filtered)" `tar.gz` file for the Fresh 68k PBMCs (Donor A) data set from the 10x Genomics website. We also downloaded the file `68k_pbmc_barcodes_annotation.tsv` from `https://github.com/10XGenomics/single-cell-3prime-paper`, which contained predicted cell-type assignments from the analysis of [35]. After filtering out genes that were not expressed in any of the cells, $\mathbf{Y}$ was a 20,387 $\times$ 68,579 matrix of UMI counts, and 97.3% of the UMI counts were zero. In plots visualizing $\mathbf{V}$, we defined "T cells" as all cells labeled as one of "CD4+ T Helper2", "CD4+/CD25 T Reg", "CD4+/CD45RA+/CD25- Naive T", "CD4+/CD45RO+ Memory" or "CD8+/CD45RA+ Naive Cytotoxic."

**3.3. Methods compared.** We compared three methods for fitting GLM-PCA models to simulated and scRNA-seq data: the AvaGrad method implemented in the `glmpca` R package [29] (version 0.2.2.9000); the iterative reweighted SVD (IRSVD) method, building on methods for "weighted SVD" [24], implemented in the `scGBM` R package [19] (version 1.0.3); and the alternating Poisson regression (APR) described in this paper and implemented in the `fastglmpca` R package (version 0.1-5). Since a key feature of the APR method is the straightforward parallelization of the updates, we compared two variants of `fastglmpca`, with and without parallelization.

Note that we did not compare the "Fisher scoring" method (`optimizer = "fisher"`) implemented in `glmpca` [27]. We found this method to be unstable—indeed this instability is documented in the R package—and others have found that generally it did not work as well as the AvaGrad method, so we did not include this method in our comparisons.

For AvaGrad, we called the `glmpca()` function from the `glmpca` package with `fam = "poi"`, `minibatch = "stochastic"` and `optimizer = "avagrad"`. For IRSVD, we called the `gbm.sc()` function from the `scGBM` package. For APR, we first called `init_glmpca_pois()` to initialize the parameter estimates, then we called `fit_glmpca_pos()` to perform the updates. For the parallel (multithreaded) variant of APR, we used 28 threads. For the DAAREM-accelerated variant of APR, we used again 28 threads, and performed the optimization in two stages: we performed an initial round of 100 updates without DAAREM, then we called `fit_glmpca_pois()` a second time with `daarem = TRUE` and `orthonormalize = FALSE`. We used version 0.7 of the `daarem` R package, setting the monotonicity tolerance (`mon.tol`) to 0.05. All other DAAREM tuning parameters were kept at their defaults. For all these calls, the remaining tuning parameters were kept at the software defaults except when modifying the termination criteria so as to force the methods to continue optimizing longer.

Unlike the other methods, the AvaGrad method is a stochastic optimization method [3, 6] and its performance can be sensitive to the choice of learning rate (despite the fact that the AvaGrad method was intended to reduce this sensitivity [23]). In particular, we found the algorithm could be unstable at higher learning rates (indeed, the "maxTry" option in `glmpca` is set by default to 10 to automatically restart the optimization in the case of numerical divergence). We experimented with different learning rates, and found that `lr = 1e-4` worked well, although it is possible that other learnings rates could have worked better.

The three software implementations included a single row covariate ($n_x = 1$) and a single column covariate ($n_z = 1$) in the model (1) by default, and we kept this default setting. These covariates were defined identically in all implementations as follows. The row covariate was a vector of ones, $x_{i1} = 1$, with fixed coefficients $b_{j1} = \log(\sum_{i=1}^{n} y_{ij}/n)$. The column covariate was also a vector of ones, $z_{j1} = 1$, with coefficients, $w_{j1}$, to be estimated. When $\mathbf{Y}$ is a matrix of UMI counts with rows $i$ corresponding to genes and columns $j$ corresponding to cells, this row covariate defines a "size factor" [1] for each cell $j$ as the mean UMI count per gene, and this column covariate encodes a baseline expression level for each gene $i$.

For fair comparison, we provided the same initial estimates of $\mathbf{U}$ and $\mathbf{V}$ to `glmpca` and `fastglmpca`. The entries of $\mathbf{U}$ and $\mathbf{V}$ were initialized to small values by randomly drawing from the normal distribution with mean zero and standard deviation $10^{-5}/K$.

The `scGBM` interface did not allow the initial estimates of $\mathbf{U}$ and $\mathbf{V}$ to be specified. Instead, the `gbm.sc()` function computes initial estimates based on Pearson residuals obtained by fitting a simple rank-2 GLM-PCA model to the data [20]. In practice, we have found that this initialization is better than our simple random initializations, but the downside is that it involves memory-intensive computations.

With the `minibatch = "stochastic"` setting, `glmpca()` does not output the exact log-likelihood achieved at each iteration. Therefore, we modified the code slightly. These additional log-likelihood calculations have a neglible impact on the overall runtime. The modified code is available at `https://github.com/eweine/glmpca`.

**3.4. Evaluation of clustering performance in 95k PBMC data set.** To cluster the cells, we ran the `hclust` function from the R package `fastcluster` [18], with `method = "ward"` [31], on the Euclidean distances computed from a scaled projection of the cells, $\mathbf{V\Sigma}$. This produced a hierarchical clustering of the cells. Then we used the `cuttree` function in R to recover 10 clusters from this hierarchical clustering. (We chose 10 clusters because the 95k PBMC data set is a mixture of 10 FACS-purified data sets.) Following [25], we computed the adjusted Rand index (ARI) and normalized mutual information (NMI) [30] to measure the similarity between the estimated clusters and the FACS cell-type labels. These calculations were performed using the `aricode` R package [8].

We repeated this clustering analysis at each iteration for each of the GLM-PCA methods compared.

**3.5. Computing environment.** All analyses were performed in R 3.6.1 [22] linked to the OpenBLAS 0.2.19 optimized numerical libraries on Linux machines (Scientific Linux 7.4) with Intel Xeon E5-2680v4 ("Broadwell") processors. Up to 56 GB of memory was needed to run `glmpca` and `fastglmpca` on the largest data set (68k PBMC). The `scGBM` computations were more memory-intensive, and needed up to 250 GB of memory on the largest data set. For the parallel variant of `fastglmpca`, we used 28 cores.

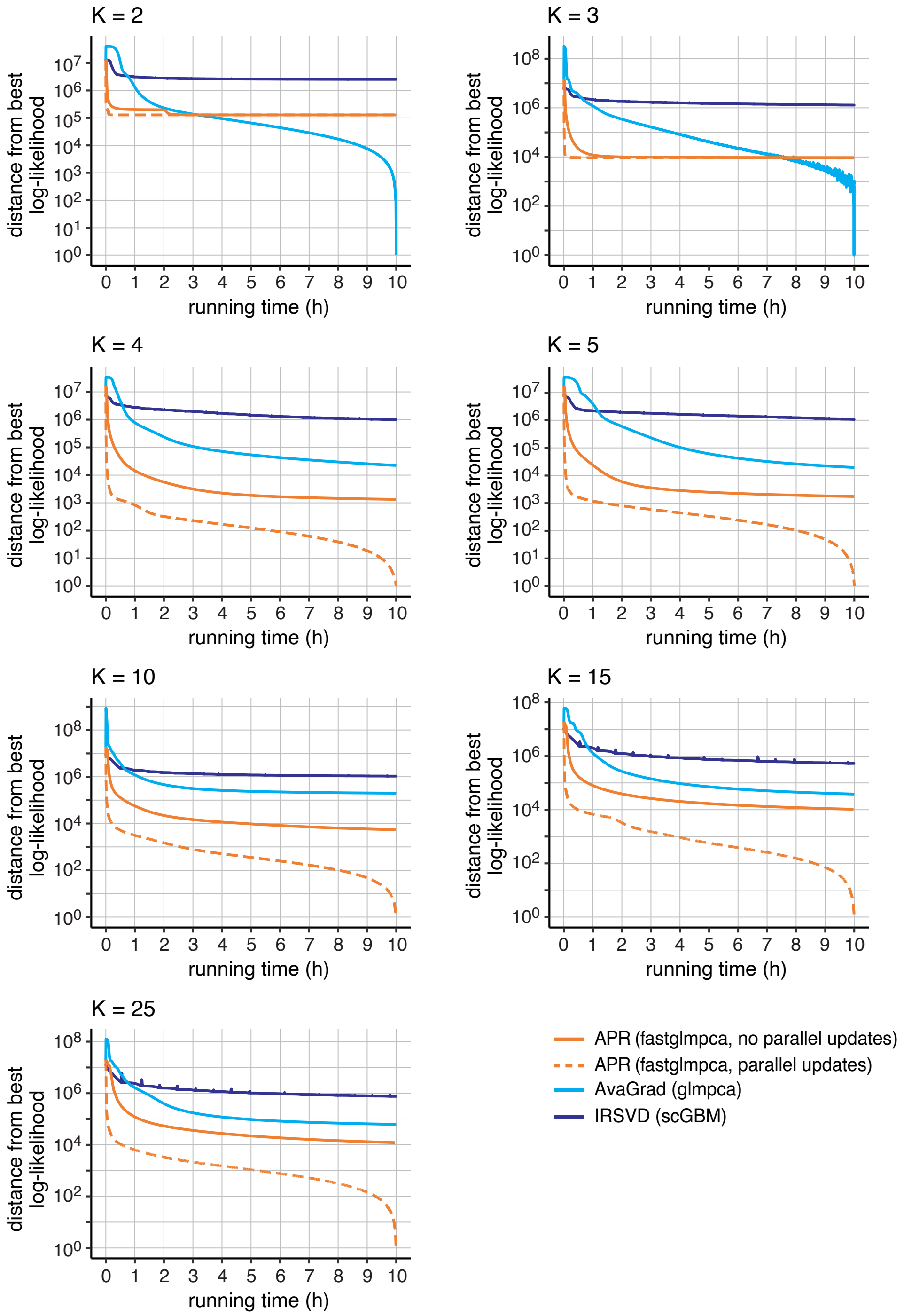


Supplementary Figure 1. Improvement in GLM-PCA model fits over time for the mouse epithelial airway data. Log-likelihoods are shown relative to the best log-likelihood recovered among the four methods compared (including the two APR variants). The Y axis has a log scale, and log-likelihood differences less than 1 are shown as 1.

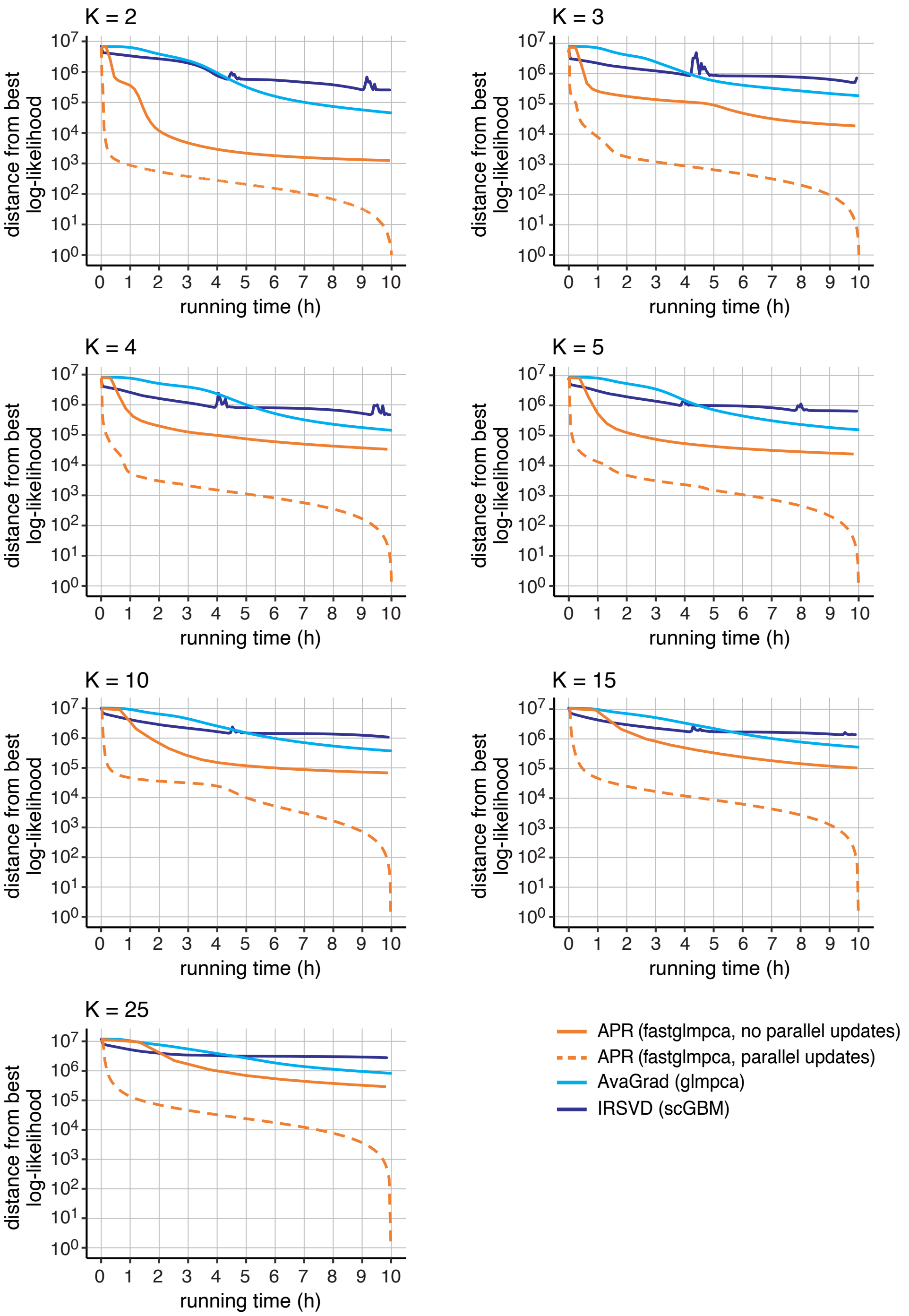


Supplementary Figure 2. Improvement in GLM-PCA model fits over time for the 68k PBMC data. Log-likelihoods are shown relative to the best log-likelihood recovered among the four methods compared (including the two APR variants). The Y axis has a log scale, and log-likelihood differences less than 1 are shown as 1.

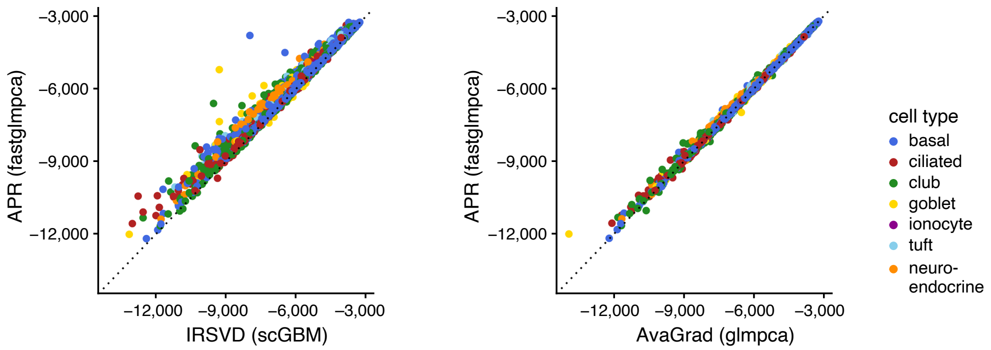


Supplementary Figure 3. Assessment of fit of the 7,193 transcriptome profiles in the epithelial airway data set cells). The X and Y axes show the log-likelihoods obtained by running the different optimization methods for about 10 hours, with $K = 10$. The `fastglmpca` result was obtained using parallel updates. Colors depict the labeling obtained by running a community detection algorithm on the scRNA-seq data [17].

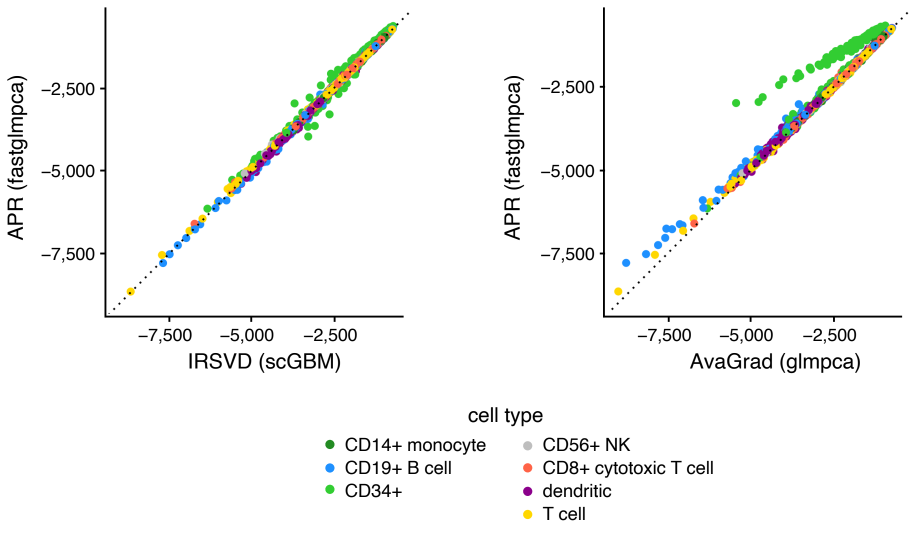


Supplementary Figure 4. Assessment of fit of the 68,579 transcriptome profiles in the 68k PBMC data set. The X and Y axes show the log-likelihoods obtained by running the different optimization methods for about 10 hours, with $K = 10$. The `fastglmpca` result was obtained using parallel updates. Colors depict the predicted cell-type annotations from [35].

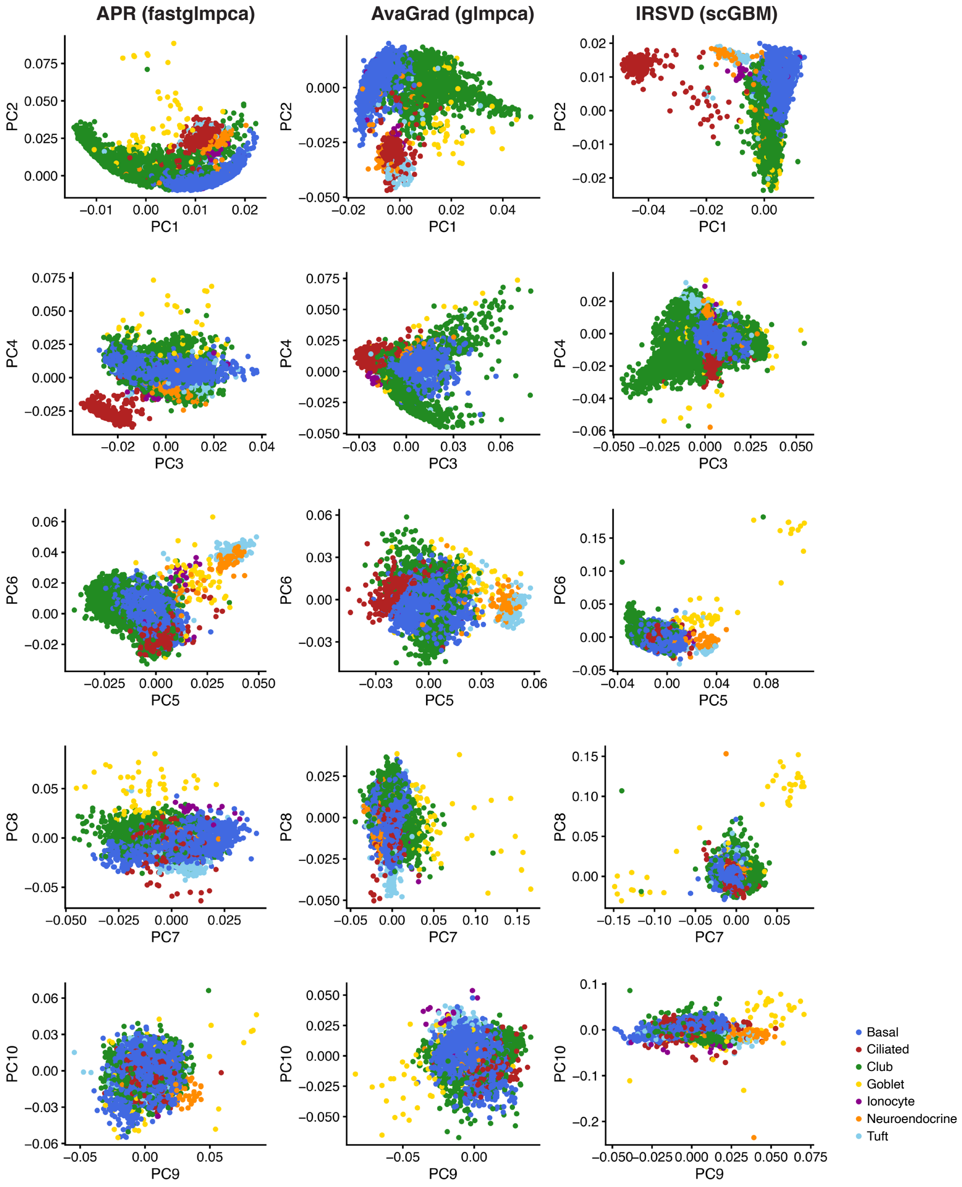


SUPPLEMENTARY FIGURE 5. The **V** matrix returned by different optimization methods when applied to the same mouse epithelial airway data set after running the optimization for about 10 hours ($K = 10$). The `fastglmpca` result was obtained using parallel updates. Colors depict the labeling obtained by running a community detection algorithm on the scRNA-seq data [17].

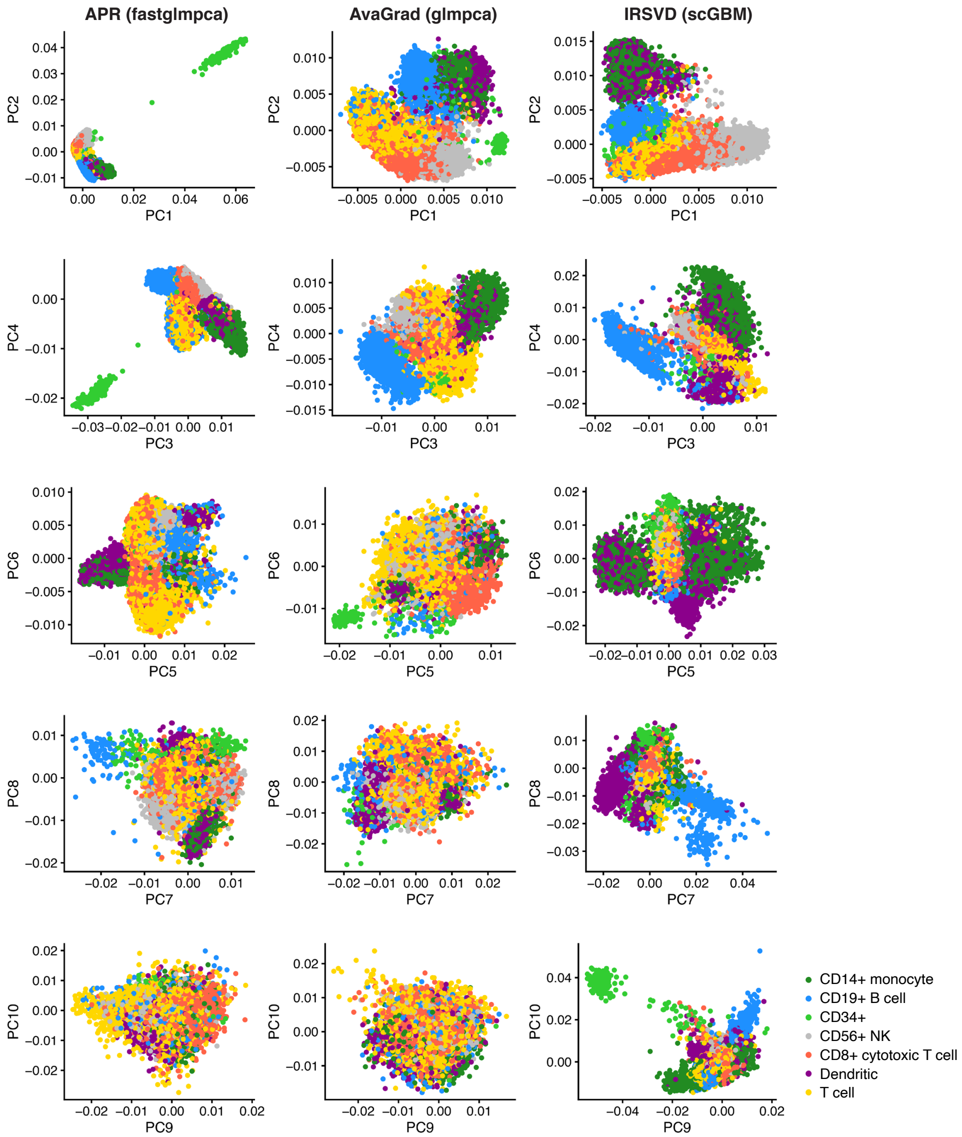


Supplementary Figure 6. The $\mathbf{V}$ matrix returned by different optimization methods when applied to the same 68k PBMC data set after running the optimization for about 10 hours ($K = 10$). The `fastglmpca` result was obtained using parallel updates. Colors depict the predicted cell-type annotations from [35].

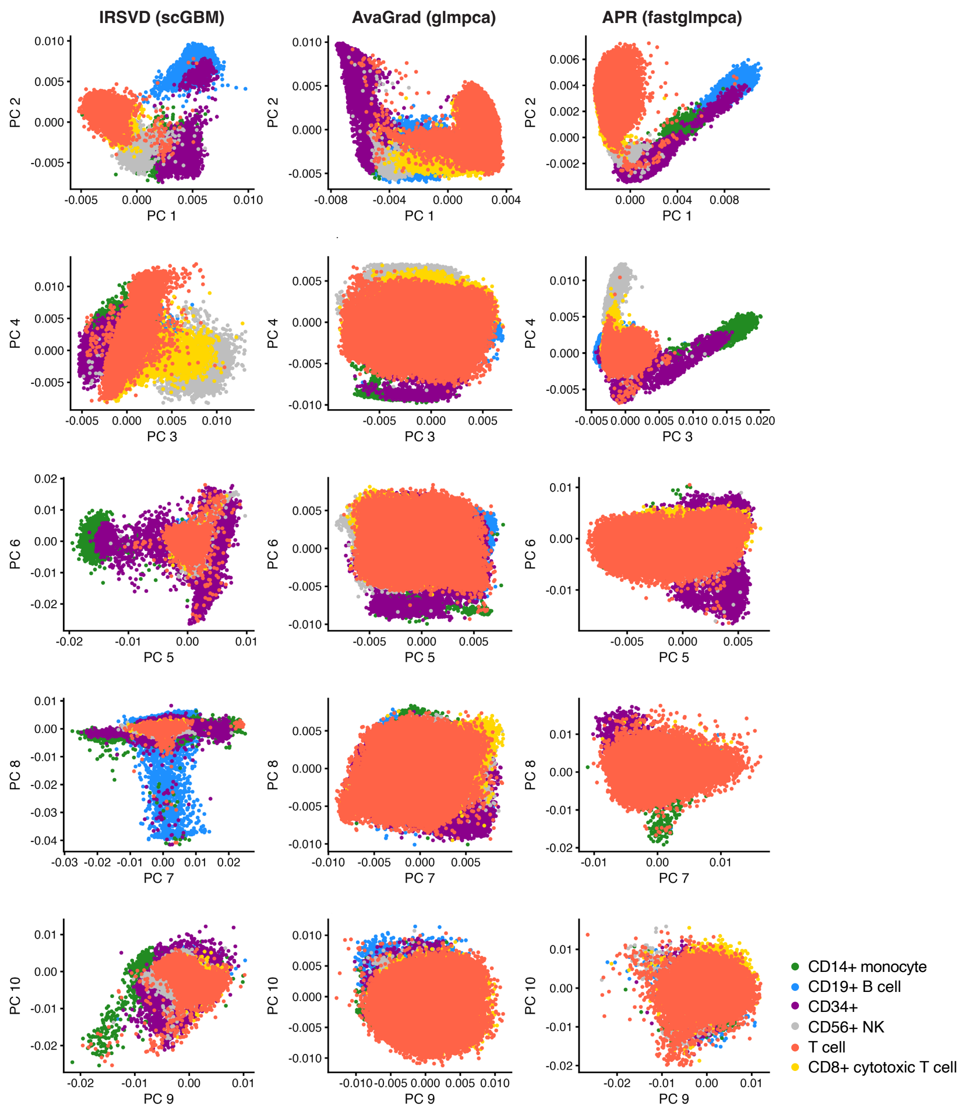


Supplementary Figure 7. The **V** matrix returned by different optimization methods when applied to the same "purified" PBMC data set after running the optimization for about 10 hours on the epithelial airway data ($K = 10$). The `fastglmpca` result was obtained using parallel updates. Colors depict the sorted cell populations. "T cells" combines all sorted T cell populations other than CD8+ cytotoxic T cells.)

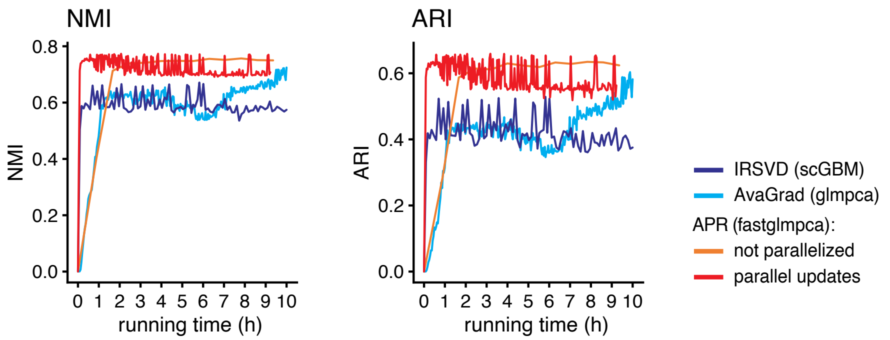


Supplementary Figure 8. Change in the clustering results over time for the 95k PBMC data. The clusters were compared against the cell population labels obtained by FACS; the similarity between the estimated clusters and "true clusters" (sorted cell populations) was measured using the normalized mutual information (left) and the adjusted Rand index (right).

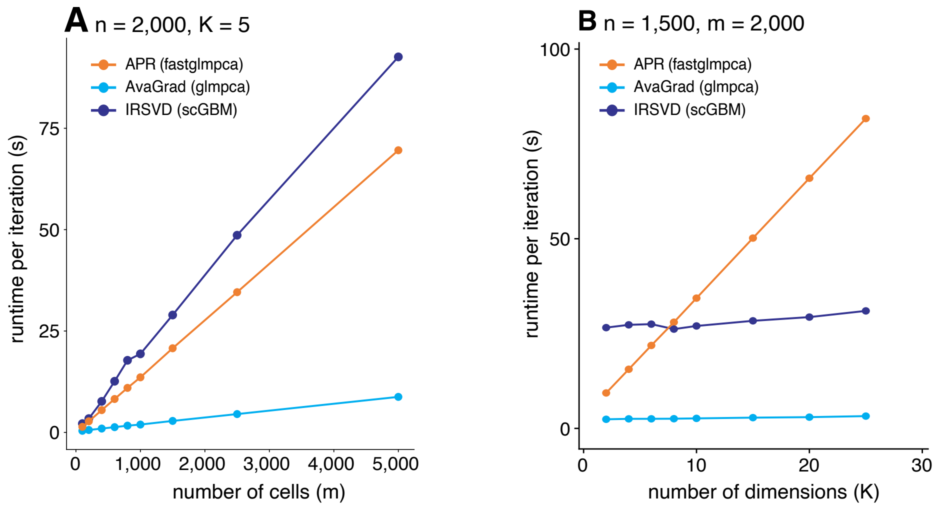


Supplementary Figure 9. Per-iteration runtimes. Panel A compares per-iteration running times for 2,000 $\times$ $m$ data matrices $\mathbf{Y}$ in which $m$ was varied, with $K = 5$. Panel B compares per-iteration running times for 1,500 $\times$ 2,000 data matrices $\mathbf{Y}$ in which $K$ was varied. Running times are averages of 100 independent runs (column A) and 50 independent runs (column B), with 10 updates performed in each run. The APR updates were performed in `fastglmpca` without parallel computation. For AvaGrad in `glmpca`, stochastic gradients were computed using 10% of the cells (mini-batches of size $m/10$).
